# Deceptive combined effects of short allele dominance and stuttering: an example with *Ixodes scapularis*, the main vector of Lyme disease in the U.S.A.

**DOI:** 10.1101/622373

**Authors:** Thierry De Meeûs, Cynthia T. Chan, John M. Ludwig, Jean I. Tsao, Jaymin Patel, Jigar Bhagatwala, Lorenza Beati

## Abstract

Null alleles, short allele dominance (SAD), and stuttering increase the perceived relative inbreeding of individuals and subpopulations as measured by Wright’s *F*_IS_ and *F*_ST_. Ascertainment bias, due to such amplifying problems are usually caused by inaccurate primer design (if developed from a different species or a distant population), poor DNA quality, low DNA concentration, or a combination of some or all these sources of inaccuracy. When combined, these issues can increase the correlation between polymorphism at concerned loci and, consequently, of linkage disequilibrium (LD) between those. In this note, we studied an original microsatellite data set generated by analyzing nine loci in *Ixodes scapularis* ticks from the eastern U.S.A. To detect null alleles and SAD we used correlation methods and variation measures. To detect stuttering, we evaluated heterozygote deficit between alleles displaying a single repeat difference. We demonstrated that an important proportion of loci affected by amplification problems (one with null alleles, two with SAD and three with stuttering) lead to highly significant heterozygote deficits (*F*_IS_=0.1, *p*-value<0.0001). This occurred together with an important proportion (22%) of pairs of loci in significant LD, two of which were still significant after a false discovery rate (FDR) correction, and some variation in the measurement of population subdivision across loci (Wright’s *F*_ST_). This suggested a strong Wahlund effect and/or selection at several loci. By finding small peaks corresponding to previously disregarded larger alleles in some homozygous profiles for loci with SAD and by pooling alleles close in size for loci with stuttering, we generated an amended dataset. Except for one locus with null alleles and another still displaying a modest SAD, the analyses of the corrected dataset revealed a significant excess of heterozygotes (*F*_IS_=-0.07 as expected in dioecious and strongly subdivided populations, with a more reasonable proportion (19%) of pairs of loci characterized by significant LD, none of which stayed significant after the FDR procedure. Strong subdivision was also confirmed by the standardized *F*_ST_’ corrected for null alleles (*F*_ST_’=0.19) and small effective subpopulation sizes (*N*_*e*_=7).

## Introduction

Null alleles, short allele dominance (SAD) and stuttering are frequent consequences of poor PCR amplifications, in particular for microsatellite markers. Amplification problems usually arise when primers are designed by using DNA of a different species or a distant population, when DNA is degraded, at too low of a concentration (Chapuis and Estoup, 2007), or any combination of these listed causes.

Null alleles occur when a mutation on the flanking sequence of the targeted locus affects the hybridization of the corresponding primer, resulting in amplification failure. Heterozygous individuals with one null allele falsely appear to be homozygous, while homozygous individuals for null alleles are considered to be missing data.

SAD, also called large allele dropout (Van Oosterhout et al., 2004), known from minisatellite markers, was also discovered to occur in microsatellite loci (Wattier et al., 1998). In heterozygous samples, longer alleles are less successfully amplified than shorter alleles through competition for Taq polymerase. This can lead to misinterpreting heterozygous individuals as homozygous for the shortest allele (De Meeûs et al., 2007).

Stuttering is the result of inaccurate PCR amplification through Taq slippage of a specific DNA strand. This generates several PCR products that differ from each other by one repeat and can cause difficulties when discriminating between fake and true homozygotes, such as heterozygous individuals for dinucleotide microsatellite allele sequences with a single repeat difference.

Allelic dropout is akin to SAD, but occurs randomly to any allele irrespective of its size, and can affect both alleles of heterozygous individuals.

The consequence of these issues is a homozygous excess when compared to the expected Castle-Weinberg proportions (Castle, 1903; Weinberg, 1908) (classically known as Hardy-Weinberg, however, more accurately depicted as Castle-Weinberg; because the former was the first to derive it for two equifrequent alleles in 1903 and the latter generalized the concept in January 1908, prior to Hardy in April 1908 (Hardy, 1908)), as measured by Wright’s *F*_IS_ (Wright, 1965). These problems, like all others associated with amplification, are locus specific (Van Oosterhout et al., 2004; De Meeûs et al., 2007; De Meeûs, 2018a) and thus lead to locus specific variation (namely, an increase) of *F*_IS_. A less well known, though well documented (Chapuis and Estoup, 2007; Séré et al., 2017b; Manangwa et al., 2019) effect of such amplification problems consists of an increase of Wright’s *F*_ST_ (Wright, 1965) that is commonly used to measure the degree of genetic differentiation between subpopulations.

While an analytical cure exists for null alleles (Chapuis and Estoup, 2007; Séré et al., 2017a), such remediation is unavailable for SAD and stuttering. To the best of our knowledge, the impact of amplification problems on linkage disequilibrium (LD) between locus pairs has yet to be investigated. Problems with amplification can be expected to more commonly occur in individuals that display some kind of deviating DNA in terms of quantity or quality: flanking sequences that have accumulated mutations, samples containing weak DNA concentration, badly preserved DNA extracts, or a combination of these different problems. When combined, the effect of the occurrence of null alleles, SAD, and stuttering may artificially generate a positive correlation between allele occurrences at affected loci and then increase the perceived LD between them.

In this note, we utilize an original data set generated through the analysis of nine microsatellite loci in *Ixodes scapularis*, sampled across the eastern U.S. to show that the combined effect of SAD, stuttering, and null alleles can induce an increase in the number of locus pairs in significant LD. We then propose and test an efficient way to amend such data.

## Material and Methods

### Sampling and DNA extraction

Larvae, nymphs, and adults of *I. scapularis* were sampled indiscriminately from different sites across the eastern U.S. on different occasions, extending from November 2001 to May 2014, by means of dragging and flagging the vegetation (Figure 1 and Table 1) (Rulison et al., 2013).

**Table 1.** State, site, GPS coordinates (decimal degrees), 12S clade membership, date of sampling, corresponding cohort membership and size of *Ixodes scapularis* adult subsamples (*N*) from the eastern U.S.A.

| State | Site | Latitude | Longitude | Clade | Date | Cohort | *N* |
| --- | --- | --- | --- | --- | --- | --- | --- |
| Alabama | AL1 | 33.24 | -86.13 | AMI | 2011 Jan | C7 | 21 |
|  | AL1 | 33.24 | -86.13 | SOI | 2011 Jan | C7 | 6 |
|  | AL1 | 33.24 | -86.13 | AMII | 2011 Jan | C7 | 18 |
|  | AL1 | 33.24 | -86.13 | SOII | 2011 Jan | C7 | 6 |
|  | AL2 | 32.95 | -87.14 | AMI | 2012 Dec | C9 | 3 |
|  | AL2 | 32.95 | -87.14 | AMII | 2012 Dec | C9 | 2 |
| Connecticut | CT1 | 41.35 | -72.76 | AMI | 2001 Nov | C1 | 3 |
|  | CT1 | 41.35 | -72.76 | AMI | 2003 Jun | C2 | 21 |
| Florida | FL1 | 30.65 | -84.21 | SOII | 2012 Dec | C9 | 4 |
|  | FL2 | 30.06 | -81.37 | AMI | 2011 Jan | C7 | 2 |
|  | FL2 | 30.06 | -81.37 | AMII | 2011 Jan | C7 | 3 |
|  | FL2 | 30.06 | -81.37 | SOI | 2011 Jan | C7 | 1 |
|  | FL2 | 30.06 | -81.37 | SOII | 2011 Jan | C7 | 4 |
| Georgia | GA1 | 32.45 | -81.78 | AMI | 2009 Dec | C5 | 11 |
|  | GA1 | 32.45 | -81.78 | SOI | 2009 Dec | C5 | 5 |
|  | GA1 | 32.45 | -81.78 | SOII | 2009 Dec | C5 | 17 |
|  | GA1 | 32.45 | -81.78 | SOIII | 2009 Dec | C5 | 1 |
| Iowa | IA1 | 42.67 | -91.59 | AMI | 2007 May | C3 | 3 |
| Louisiana | LA1 | 30.94 | -91.46 | AMI | 2012 Dec | C9 | 3 |
|  | LA1 | 30.94 | -91.46 | AMII | 2012 Dec | C9 | 2 |
|  | LA1 | 30.94 | -91.46 | SOIII | 2012 Dec | C9 | 1 |
|  | LA1 | 30.94 | -91.46 | SOIV | 2012 Dec | C9 | 2 |
| Massachusetts | MA1 | 41.71 | -69.92 | AMI | 2010 May | C5 | 6 |
| Maryland | MD1 | 39.29 | -76.88 | AMI | 2010 Oct | C7 | 11 |
| North-Carolina | NC1 | 33.91 | -78.39 | AMI | 2009 Jan | C4 | 22 |
|  | NC1 | 33.91 | -78.39 | SOI | 2009 Jan | C4 | 4 |
|  | NC1 | 33.91 | -78.39 | SOIII | 2009 Jan | C4 | 2 |
| New-York | NY1 | 40.76 | -72.83 | AMI | 2009 Oct | C5 | 10 |
| Pennsylvania | PA1 | 40.67 | -75.96 | AMI | 2010 Oct | C7 | 5 |
|  | PA2 | 41.06 | -79.48 | AMI | 2014 May | C10 | 33 |
| South-Carolina | SC1 | 33.33 | -81.66 | AMI | 2011 Apr | C7 | 8 |
|  | SC1 | 33.33 | -81.66 | AMII | 2011 Apr | C7 | 1 |
|  | SC1 | 33.33 | -81.66 | SOI | 2011 Apr | C7 | 3 |
|  | SC1 | 33.33 | -81.66 | SOII | 2011 Apr | C7 | 1 |
|  | SC1 | 33.33 | -81.66 | SOIII | 2011 Apr | C7 | 1 |
|  | SC2 | 33.23 | -81.73 | AMI | 2010 Dec | C7 | 4 |
|  | SC2 | 33.23 | -81.73 | AMII | 2010 Dec | C7 | 8 |
|  | SC2 | 33.23 | -81.73 | SOI | 2010 Dec | C7 | 4 |
|  | SC2 | 33.23 | -81.73 | SOII | 2010 Dec | C7 | 10 |
|  | SC3 | 33.15 | -81.61 | SOII | 2010 Dec | C7 | 2 |
|  | SC5 | 34.29 | -79.87 | AMI | 2012 Dec | C9 | 3 |
|  | SC5 | 34.29 | -79.87 | SOII | 2012 Dec | C9 | 2 |
| Tennessee | TN1 | 35.37 | -86.07 | AMI | 2010 Dec | C7 | 23 |
| Texas | TX1 | 31.80 | -96.23 | AMI | 2012 Dec | C9 | 5 |
|  | TX1 | 31.80 | -96.23 | SOI | 2012 Dec | C9 | 2 |
|  | TX1 | 31.80 | -96.23 | SOIV | 2012 Dec | C9 | 3 |
|  | TX2 | 30.24 | -100.72 | AMI | 2012 Dec | C9 | 1 |
|  | TX2 | 30.24 | -100.72 | SOIV | 2012 Dec | C9 | 5 |
|  | TX3 | 31.91 | -95.9 | AMI | 2012 Dec | C9 | 3 |
|  | TX3 | 31.91 | -95.9 | SOI | 2012 Dec | C9 | 2 |
|  | TX3 | 31.91 | -95.9 | SOIV | 2012 Dec | C9 | 6 |
|  | TX4 | 31.59 | -95.61 | AMI | 2012 Dec | C9 | 2 |
|  | TX4 | 31.59 | -95.61 | SOI | 2012 Dec | C9 | 2 |
| Virginia | VA1 | 38.29 | -78.29 | AMI | 2010 Oct | C7 | 7 |
|  | VA2 | 37.32 | -80.73 | AMI | 2012 Dec | C9 | 4 |
| Wisconsin | WI1 | 43.95 | -90.70 | AMI | 2011 Oct | C8 | 8 |
|  | WI1 | 43.95 | -90.70 | AMI | 2012 Oct | C9 | 4 |
|  | WI2 | 44.04 | -90.65 | AMI | 2011 Oct | C8 | 8 |
|  | WI2 | 44.04 | -90.65 | AMI | 2012 Oct | C9 | 12 |
|  | WI3 | 44.02 | -90.64 | AMI | 2011 Oct | C8 | 8 |
|  | WI3 | 44.02 | -90.64 | AMI | 2012 Oct | C9 | 3 |

**Figure 1.**
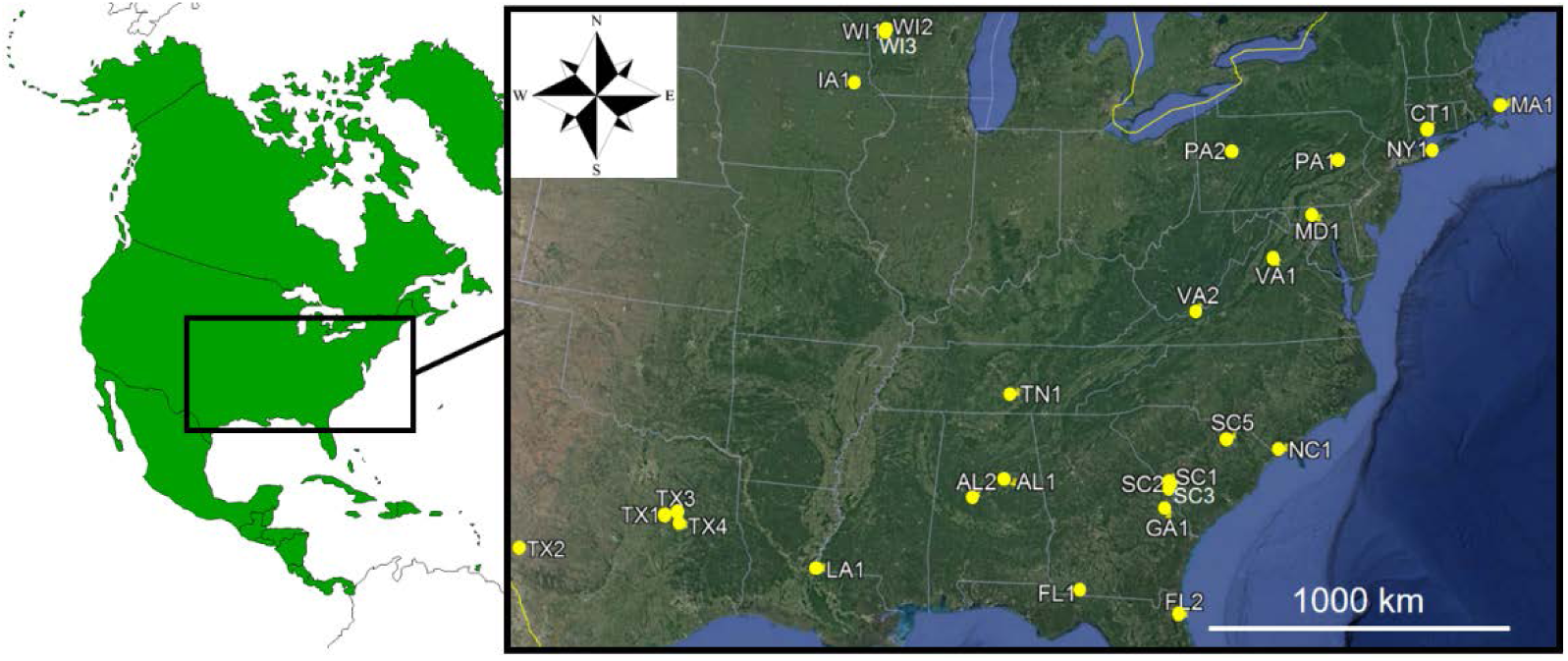
Sampling sites for *Ixodes scapularis* from the eastern U.S.A. (State codes as in Table 1).

Gravid females fall on the ground where they lay thousands of eggs at the same place that hatch as weakly mobile larvae. Larval collections can thus be composed of clusters of thousands of sisters and brothers within the same subsample (Kempf et al., 2011). Consequently, to avoid possible Wahlund effects, where a heterozygote deficit results from the admixture of individuals from genetically distant subpopulations (e.g. see (De Meeûs, 2018a)) (here families), we removed immature stages from the present study. The remaining 387 adult ticks were subdivided into cohorts, with each cohort comprised of samples collected across two consecutive years in the fall, the following winter and spring across the tick distribution range. This subdivision was based on observations showing that northeastern adults active in Fall can undergo winter quiescence and resume activity in spring (Yuval and Spielman, 1990).

Many publications have emphasized the importance of mitochondrial clades in different populations of *I. scapularis* across the U.S. (Norris et al., 1996; Qiu et al., 2002; Sakamoto et al., 2014). Thus, to account for the mitochondrial clade representation and to (again) avoid possible Wahlund effects, all ticks were assigned clades by phylogenetic analysis of their 12S rDNA gene sequences. We identified 6 main clades in our dataset, the previously identified American clade was subdivided in two lineages (AMI and AMII), and the so-called southern clade was subdivided in 4 lineages (SOI, SOII, SOIII and SOIV) (Table 1).

In conclusion, the combination of Site-Clade-Cohort data defined 61 subsamples within the 387 individual adult ticks. Overall, 35 subsamples included a small number of individuals (1-4), 12 subsamples contained at least 10 individuals and 5 subsamples contained at least 20 individuals (Table 1). Because the smallest subsamples were expected to exert a negligible weight on our analyses, they were not eliminated.

Procedures for all DNA extractions followed modified published protocols (Beati and Keirans, 2001) with a DNeasy Blood & Tissue Kit (Qiagen, Valencia, CA).

### Selection and characterization of microsatellite markers

Thirteen of the first batches of genome sequences of *I. scapularis* that were accessioned by VectorBase (www.vectorbase.org; Giraldo-Calderón *et al*. 2015) in GenBank (AC205653.1, AC205652.1, AC205650.1, AC205647.1, AC205646.1, AC205643.1, AC205642.1, AC205641.1, AC205638.1, AC205635.1, AC205634.1, AC205632.1, AC205630.1) were used to manually detect motifs with at least 6 repeats of AG, AT, CA, TA, TG, CT, GC, ACG, GTT, TTA, CAC, GAT, and AAAC. Primer pairs were selected in the flanking regions by using Oligo v.5 (Molecular Biology Insights, Colorado Springs, CO). DNA, extracted and pooled from six ticks from Connecticut, was used to test whether the selected primer sets successfully amplified fragments of the expected size. PCRs were performed using the 5-Prime Master PCR kit (5-Prime, Gaithersburg, MD) and a single touch-down amplification protocol consisting of 5 min. denaturation at 93°C; 5 cycles: 20 sec. denaturation at 93°C, 20 sec. annealing at 55°C-1.5°C/cycle, 30 sec. elongation at 72°C; 30 cycles: 20 sec. denaturation at 93°C, 25 sec. annealing at 47°C, 30 sec. elongation at 70°C; final extension at 70°C for 5 min. Amplicons were run on 4% E-gels (Life Technologies Co., Carlsbad, CA). The risk of having selected primers within repeated portions of the genome had to be considered due to the fact that large repeated genomic fragments are known to occur abundantly in *I. scapularis* (Gulia-Nuss et al., 2016). In order to confirm that the primers were amplifying the targeted loci, the amplicon of one randomly chosen tick was cloned with a TOPO-TA PCR cloning kit (Life Technologies Co, Carlsbad, CA) for each locus. Five cloned colonies were picked randomly for each tick and the insert amplified and sequenced (DNA Analysis Facility on Science Hill, Yale University). Finally, as microsatellite primers are known to amplify sometimes more than one closely related species, the same set of primers was tested on DNA samples of *Ixodes ricinus, Ixodes pacificus*, and *Ixodes persulcatus* (LB, personal collection), all taxa belonging to the *I. ricinus* complex of ticks (Keirans et al., 1999).

The primer pairs that yielded amplicons of the expected size were then used to individually amplify a subset of 67 DNA samples from ticks, representative of the distribution area of *I. scapularis* in USA, and collected by flagging or dragging in Alabama (10 ticks), Georgia (15), Connecticut (16), Massachusetts (14), New York (2), Pennsylvania (2), and South Carolina (8). For these amplifications, forward primers were labeled with fluorescent dyes (Applied Biosystems, Thermo Fisher Scientific, CA) as listed in Table 2. The amplicons were sent to the DNA Analysis Facility on Science Hill (Yale University, New Haven, CT) for genotyping. The allele peaks were scored using GeneMarker (SoftGenetics, State College, PA). All data were recorded in an Excel spreadsheet for further ease of conversion.

**Table 2.**
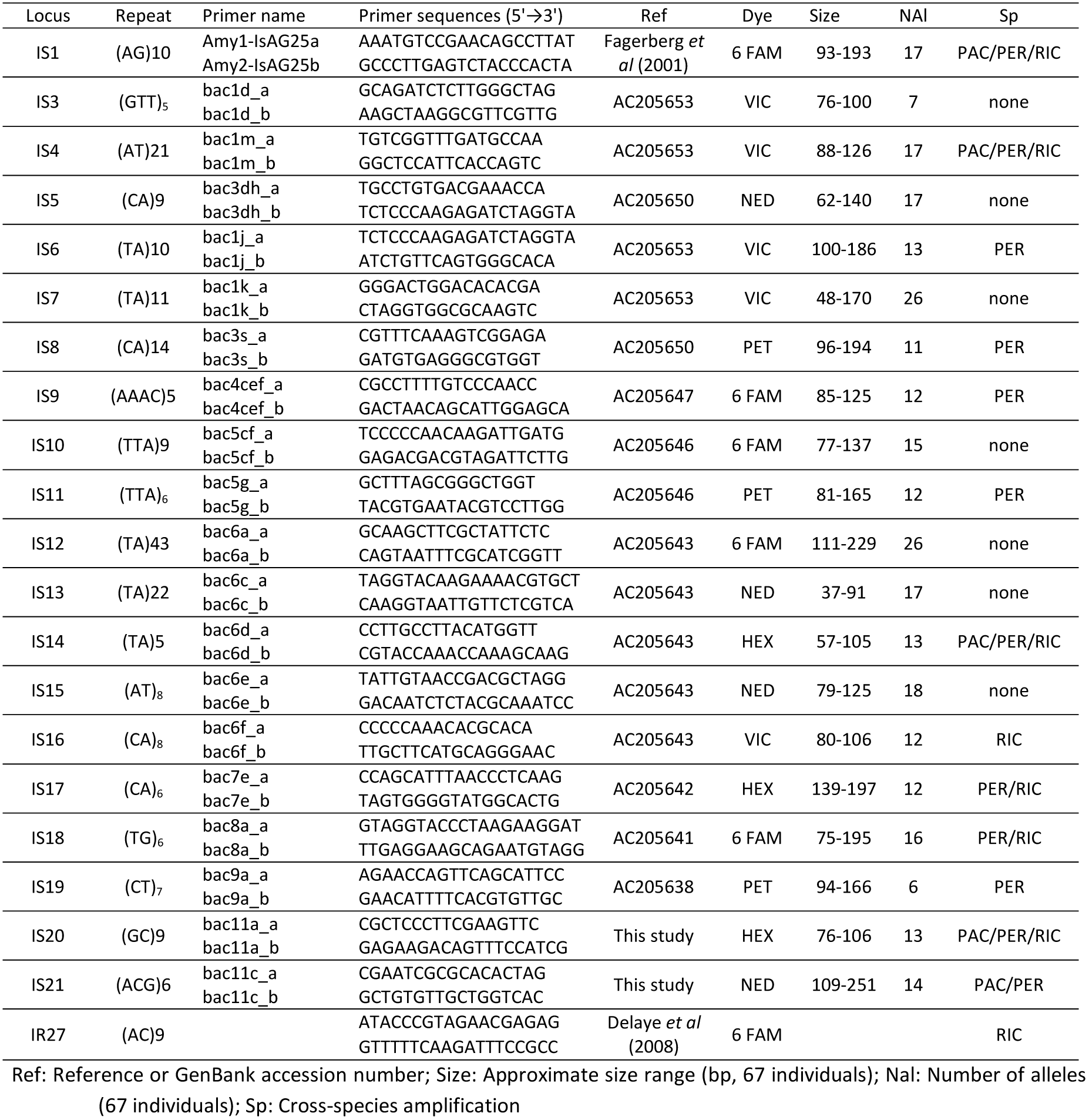
List of primer sets used or developed for this study. PAC = *Ixodes pacificus*, PER = *I. persulcatus*, RIC = *I. ricinus*.

### Genotyping

Based on their degree of polymorphism, nine microsatellite loci (IR27, IS1, IS3, IS11, IS15, IS16, IS17, IS18, and IS19) were used for genotyping at the continental scale (Table 2). Of these, IR27 (Delaye et al., 1998) and IS1 (Fagerberg et al., 2001), were drawn from previously published studies. The loci were amplified and genotyped using the procedures described above, although PCR conditions had to be slightly optimized for markers IS11 and IS15 (touchdown annealing temperature decreased from 58°C to 50°C) and IR27 (touchdown annealing temperature decreased from 56°C to 53°C) (Table 2).

### Population genetics analyses

The raw data set was coded and converted into all required formats using Create (Coombs et al., 2008).

To test for LD, we used the *G*-based test first described by Goudet et al. (Goudet et al., 1996) and adapted for contingency tables of locus pairs, with 15000 reshuffling of genotypes to get maximum precision and minimize possible *p*-values before false discovery rate corrections (see below). The *G* statistics obtained for each subsample were then summed over all subsamples to get a single statistic and hence, a single test across subsamples. This procedure was shown to be the most powerful (De Meeûs et al., 2009) and was implemented within Fstat 2.9.4 (Goudet, 2003) an updated version of the original 1995 Fstat software (Goudet, 1995). There are as many tests as locus pairs and these tests are correlated (one locus is used as many times as there is any other locus). To take into account this repetition of correlated tests, we used Benjamini and Yekutieli (BY) false discovery rate procedure (Benjamini and Yekutieli, 2001) with R version 3.5.1 (R-Core-Team, 2018). To check if some loci were involved in a significant LD pair more often than by chance, as compared to the other loci, we also undertook a Fisher exact test with Rcmdr version 2.3-1 (Fox, 2005; Fox, 2007).

For a hierarchy with three levels (individuals, subsamples, and total sample), three *F*-statistics can be defined (Wright, 1965). *F*_IS_ measures inbreeding of individuals relative to inbreeding of subsamples or relative deviation of observed genotypic proportions from local random mating proportions. *F*_ST_ measures inbreeding of subsamples relative to total inbreeding or relative inbreeding due to the subdivision of the total population into several isolated subpopulations. *F*_IT_ measures inbreeding of individuals relative to total inbreeding. Under the null hypothesis (panmixia and no subdivision), all these statistics are expected to be null. Otherwise, *F*_IS_ and *F*_IT_ can vary from -1 (one heterozygote class) to +1 (all individuals homozygous) and *F*_ST_ from 0 (all subsamples share similar allele frequencies) to +1 (all subsamples fixed for one or the other allele). These statistics were estimated with Weir and Cockerham’s unbiased estimators (Weir and Cockerham, 1984) with Fstat.

In dioecious species (like ticks), heterozygote excess occurs over all loci (e.g. (De Meeûs et al. 2007)) and is proportional to subpopulation size (*N*_*e*_=-1/(2×*F*_IS_)-*F*_IS_/(1+*F*_IS_)) (Balloux, 2004). Therefore, the finding of homozygous excesses really represents a strong deviation from random mating expectations. Technical problems, like null alleles, stuttering, SAD or allele dropouts unevenly affects some loci, producing a positive *F*_IS_ with an important variation across loci and significant outliers (De Meeûs 2018). Significant departure from 0 of these *F*-statistics was tested with 10000 randomizations of alleles between individuals within subsample (deviation from local random mating test) or of individuals between subsamples within the total sample (population subdivision test). For *F*_IS_, the statistic used was *f* (Weir and Cockerham’s *F*_IS_ estimator). To test for subdivision, we used the *G*-based test (Goudet et al. 1996) over all loci, which is the most powerful procedure when combining tests across loci (De Meeûs et al. 2009).

To compute 95% confidence intervals (95%CI) of *F*-statistics, we used the standard error of *F*_IS_ (StrdErrFIS) and *F*_ST_ (StrdErrFST) computed by jackknifing over populations, or 5000 bootstraps over loci as described elsewhere (De Meeûs et al. 2007). For jackknives, the number of usable subsamples was restricted to subsamples with at least 5 ticks (23 subsamples) (e.g. (De Meeûs, 2012) p 73).

In case of significant homozygote excess and LD we tried to discriminate demographic from technical causes with the determination key proposed by De Meeûs (De Meeûs 2018). Null alleles better explain the data if the StrdErrFIS becomes at least twice as high as StrdErrFST; *F*_IS_ and *F*_ST_ are positively correlated; and a positive correlation links *F*_IS_ and the number of missing data (putative null homozygotes). The significance of correlations was tested with a unilateral (*ρ*>0) Spearman’s rank correlation test with R. The presence of null alleles was also verified with MicroChecker v 2.2.3 (Van Oosterhout et al. 2004) and null allele frequencies estimated with Bookfield’s second method (Brookfield 1996). The adjustment between observed and expected numbers of missing data was tested with a unilateral exact binomial test in R with the alternative hypothesis being “there is not enough missing data as expected if heterozygote deficits were entirely explained by null alleles under panmixia”. MicroChecker also seeks stuttering and SAD. Stuttering is detected when MicroChecker reveals an observed proportion of heterozygous individuals for alleles with one repeat difference significantly smaller than the expected value. The presence of stuttering was detected with the graphic output of MicroChecker (we ignored the comments panel that happened to contradict the graphic in some instances). We considered that the observed deficit of heterozygous individuals for one repeat difference was a likely consequence of stuttering. Due to the small population sample sizes and bootstrapping procedure in MicroChecker, the statistical support (*p*-value) of this result was not always significant for all runs. Hence, we set the randomization at the maximum value (10000) and repeated the analysis three times to check for consistency. Stuttering was admitted when two out of three tests supported it. The occurrence of SAD was also checked with an unilateral (*ρ*<0) Spearman’s rank correlation test between allele size and *F*_IT_ as proposed by (Manangwa et al., 2019). This test is more powerful than with *F*_IS_ as was proposed earlier (Wattier et al., 1998; De Meeûs et al., 2004). If previous tests are not significant and if StrdErrFIS > StrdErrFST, then a Wahlund effect better explains the data (De Meeûs, 2018a), this is especially true in instances of significant LD (Manangwa et al., 2019). In these cases, a positive correlation between the number of times a locus is found in significant LD (NLD) and its total genetic diversity as measured by Nei’s *H*_T_ (Nei and Chesser, 1983) (Spearman’s correlation above 0.1) suggests an admixture of individuals from several subpopulations of the same species but with an important degree of genetic differentiation between admixed subpopulations (i.e. number of immigrants *N*_*e*_*m*=2). If the correlation is negative and the proportion of significant LDs is above 40%, an admixture of different strongly divergent entities (e.g. species) better explains the data (Manangwa et al., 2019). We thus undertook a bilateral Spearman’s test.

In some instances, the same null hypothesis was repeatedly tested (i.e. SAD was tested for each locus one by one). Repetition of independent tests were submitted to Benjamini and Hochberg (BH) correction (Benjamini and Hochberg, 2000) (computed with R), to check for robustness of significant *p*-values.

To our knowledge, when diagnosed, there is no analytical remedy for SAD or stuttering as there is for null alleles (Chapuis and Estoup, 2007; Séré et al., 2017a). Nevertheless, SAD can be cured by going back to the chromatograms of homozygous individuals and trying to find a micro-peak (see the Results and discussion section), with a larger size, ignored in the first reading. If enough profiles can be corrected this might salvage the incriminated locus.

Stuttering can be addressed by pooling alleles close in size. However, this procedure requires that none of the pooled allele groups is constituted of rare alleles only. Indeed, pooling rare alleles, usually found in heterozygosity with a more frequent allele, will tend to artificially generate heterozygous excesses. In order to avoid this consequence, each pooled group should contain at least one frequent allele (e.g. with *p*≥0.05).

In dioecious small populations, a heterozygote excess is expected. However, loci with null alleles may display heterozygote deficits. In such situations a bilateral test (*F*_IS_ is not different from 0) is needed and obtained as *p*_bilateral_=*p*_mini_+1-*p*_maxi_, where *p*_mini_ is the minimum unilateral *p*-value (for heterozygote deficit or excess) and *p*_max_ is the maximum one.

Finally, a more accurate estimate of *F*_ST_ can be made for datasets with null alleles after recoding missing genotypes as homozygous for allele 999 (null allele) with the ENA method as implemented in FreeNA (Chapuis and Estoup, 2007). This estimate can then be corrected for polymorphism with the formula *F*_ST_’=*F*_ST_/(1-*H*_S_) (Hedrick, 1999). This helped compute the number of immigrants as *Nm*=(1-*F*_ST_’)/(4*F*_ST_) (assuming an Island model of migration) (e.g. (De Meeûs et al., 2007)).

## Results and discussion

### Primer selection and characterization of loci

The inspection of the GenBank genomic sequences revealed the presence of 67 short tandem repeated motifs. The program Oligo v.5 did not find suitable primers for 17 of them. Of the remaining 50 primer pairs, 22 amplified the pooled DNA sample and the sizes of the amplicons were approximately as expected. The cloned amplified inserts all contained the expected microsatellite repeats and flanking regions. The 22 primer sets consistently amplified DNA from the 67 *I. scapularis* ticks and some of them also amplified DNA of the other *Ixodes* species (Table 2). Finally, based on their polymorphism and ease of interpretation, nine loci were retained for the population genetics analyses: IR27 (from (Delaye et al., 1998)), IS1, IS3, IS11, IS15, IS16, IS17, IS18, and IS19 (from the present study), with 9.8, 3.4, 1.3, 8, 8.3, 4.9, 2.8, 5.7 and 4.1 % missing genotypes (blanks) respectively.

### Raw data analyses

All data are available in the supplementary File S1.

There was a very important proportion of locus pairs in significant LD (∼22%), with a negative correlation between NLD and *H*_T_ (*ρ*=-0.04, *p*-value=0.9242). Two locus pairs remained in significant LD after BY correction: IR27-IS3 (*p*-value=0.0301) and IS11-IS16 (*p*-value=0.0451). No single locus was found more often in significant LD than the others (*p*-value=0.09).

There was a highly significant heterozygote deficit (*F*_IS_=0.111, in 95%CI=[-0.02, 0.236, *p*-value<0.0002), with a considerable variation across loci (Figure 2).

**Figure 2.**
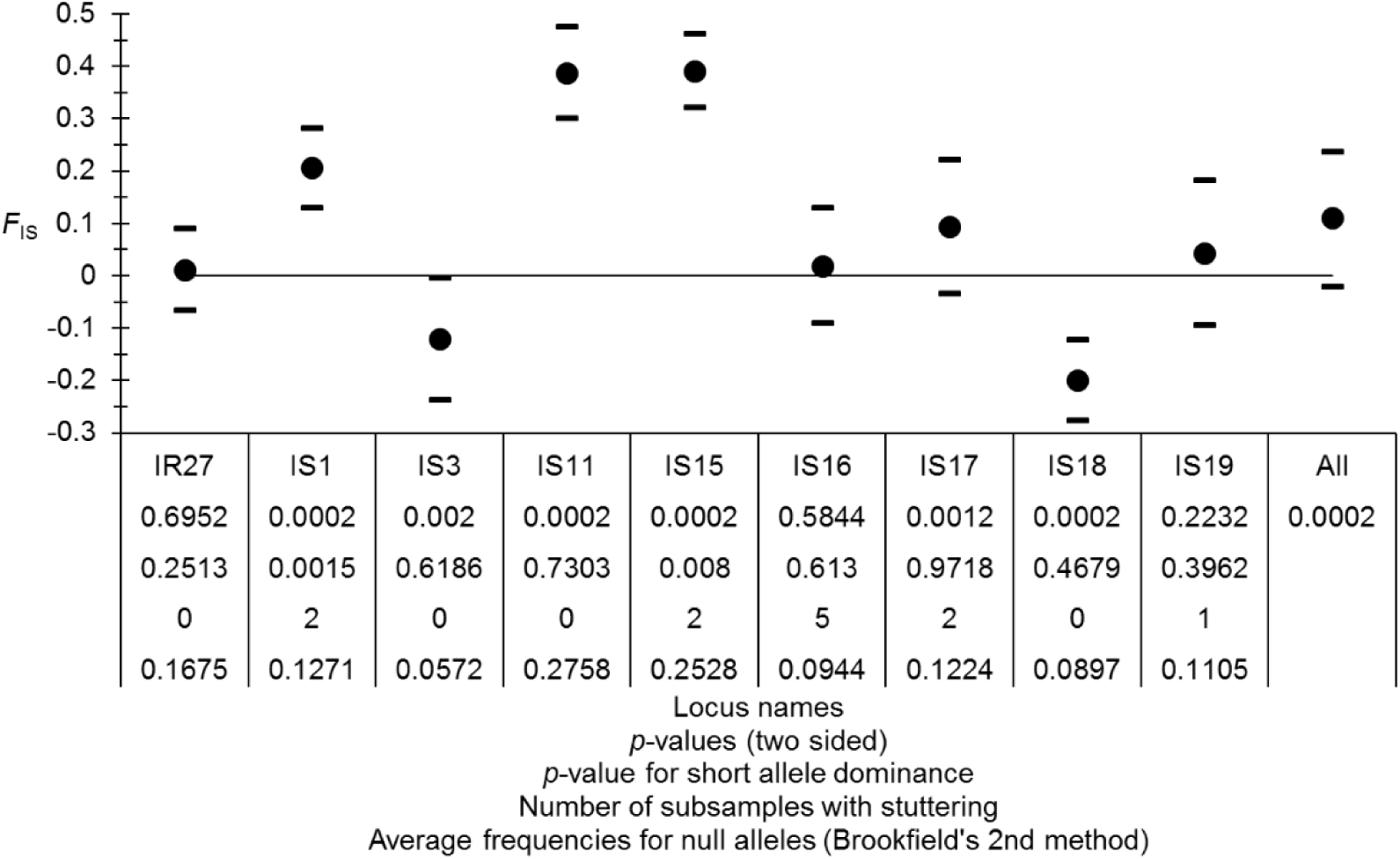
*F*_IS_ values for each locus and averaged across those (All) of *Ixodes scapularis* from the eastern U.S.A. with 95% jackknife confidence intervals over subsamples (for each locus) and bootstraps over loci (All). Results of tests for panmixia, short allele dominance, number of subsamples with stuttering for each locus, and null allele frequencies are also indicated.

StdrdErrFIS (0.07) was almost four times higher than StrdErrFST (0.019); the correlation between *F*_IS_ and *F*_ST_ was negative (*ρ*=-0.6166, *p*-value=0.962) and the correlation between *F*_IS_ and the number of blanks (missing genotypes) was positive but not significant (*ρ*=0.1833, *p*-value=0.3218). These results suggested that locus-specific effects were involved. Nevertheless, null alleles only partly explained the observed pattern at best. The substantial proportion of significant LDs suggested the existence of a possible strong Wahlund effect, though a negative correlation between NLD and *H*_T_ with less than 40% significant tests observed here would refute this interpretation (Manangwa et al., 2019).

Small subsampling due to the partitioning of the data into cohorts and 12S clades was expected to considerably lower the power of these tests, especially so after the rather stringent BY procedure for the LD test series.

Subdivision was substantial (*F*_ST_=0.101, 95%CI=[0.066, 0.136], *p*-value<0.0001), and variable across loci and subsamples (Figure 3), but not meaningfully more variable than expected (see Figure 6 in (De Meeûs, 2018b)). Some loci (IS1, IS15 and IS17) displayed very low values (Figure 3). Selection (e.g. balanced selection) might have produced the pattern observed, though evidence for this is weak (large confidence intervals).

**Figure 3.**
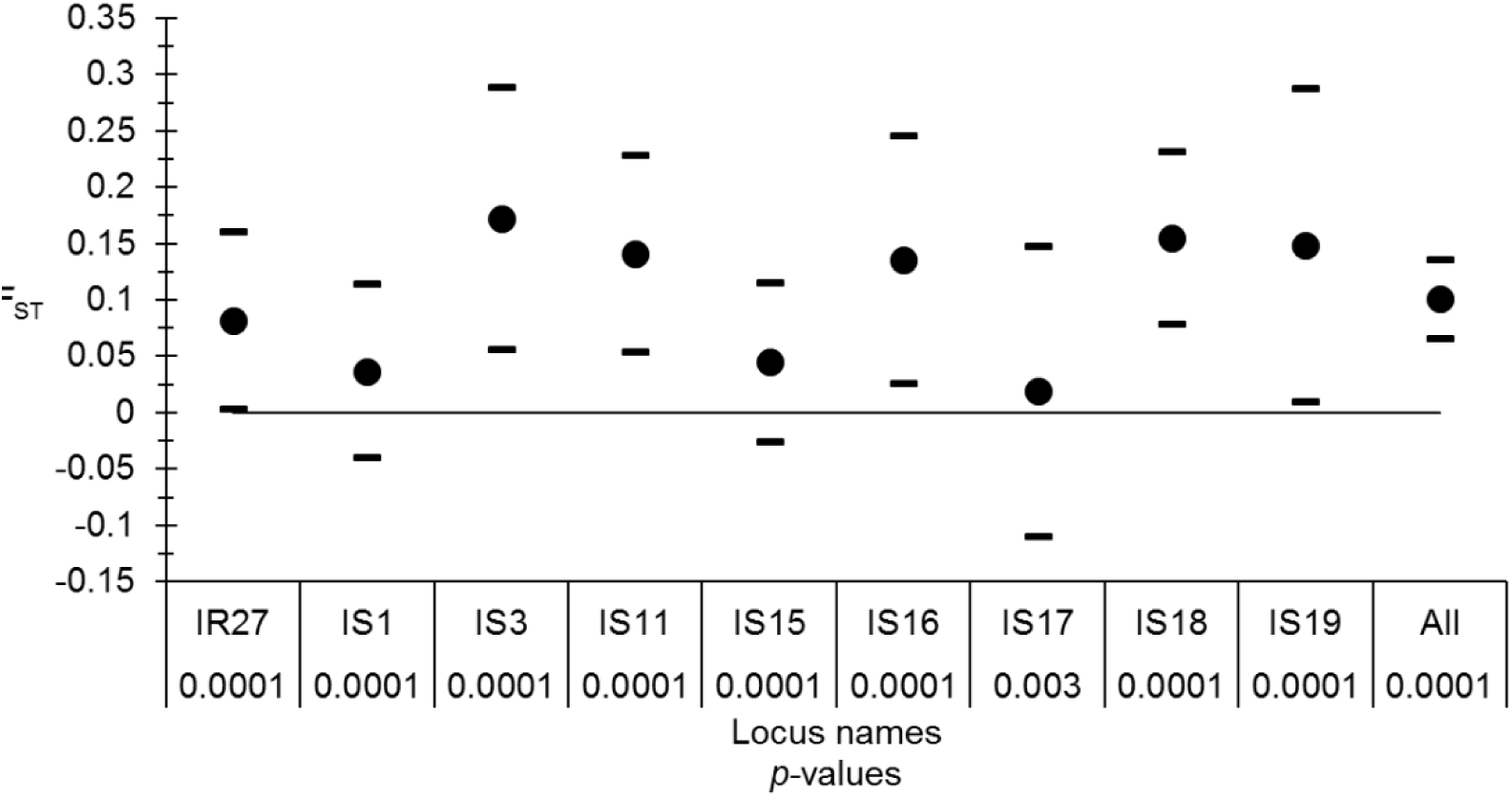
*F*_ST_ values for each locus and averaged across those (All) of *Ixodes scapularis* from the eastern U.S.A. with 95% jackknife confidence intervals over subsamples (for each locus) and bootstrap over loci (All). Results of tests for significant subdivision (*p*-value) are also indicated.

**Figure 4.**
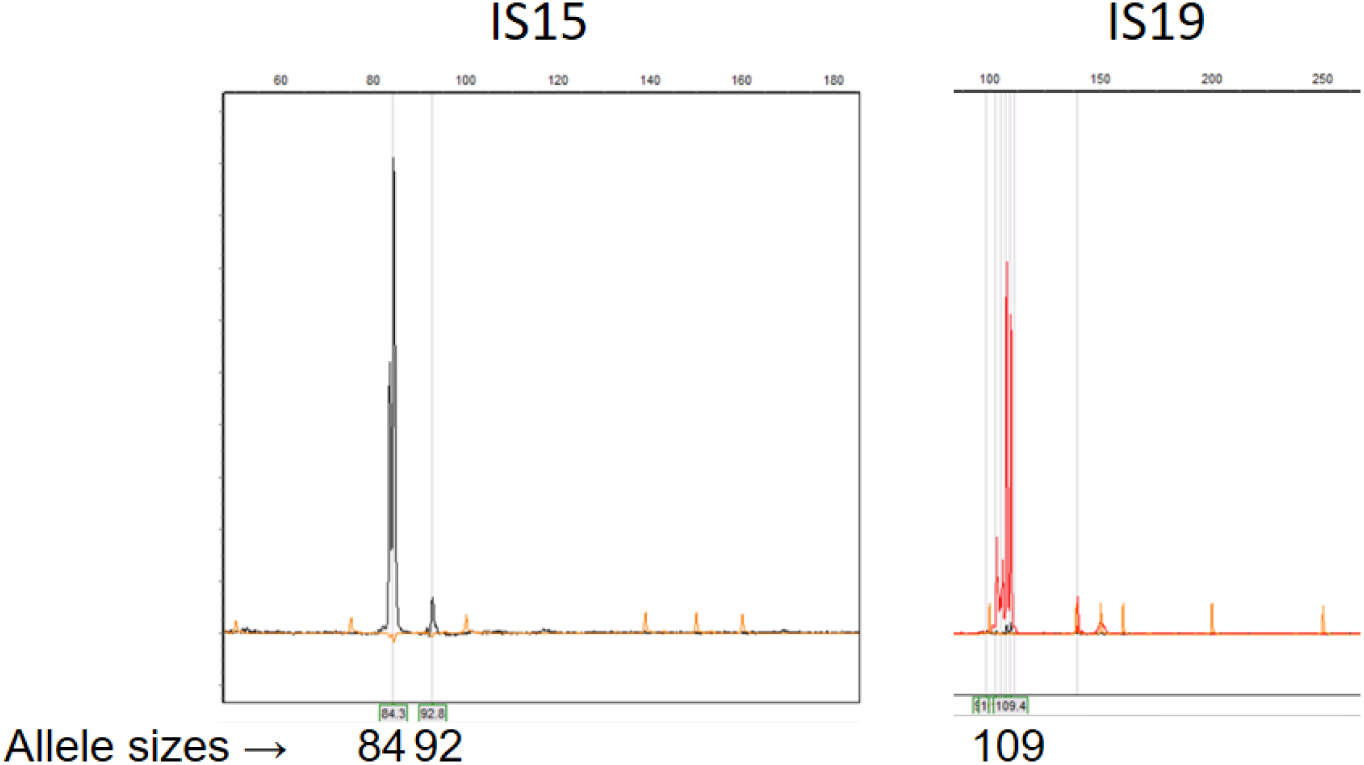
Examples of an initially dismissed micro-peak that produced SAD at locus IS15 (the peak for allele 92 appears much smaller than for allele 84) and of stuttering at locus IS19 (stutters around allele 109 hides the possible presence of allele 111).

**Figure 5.**
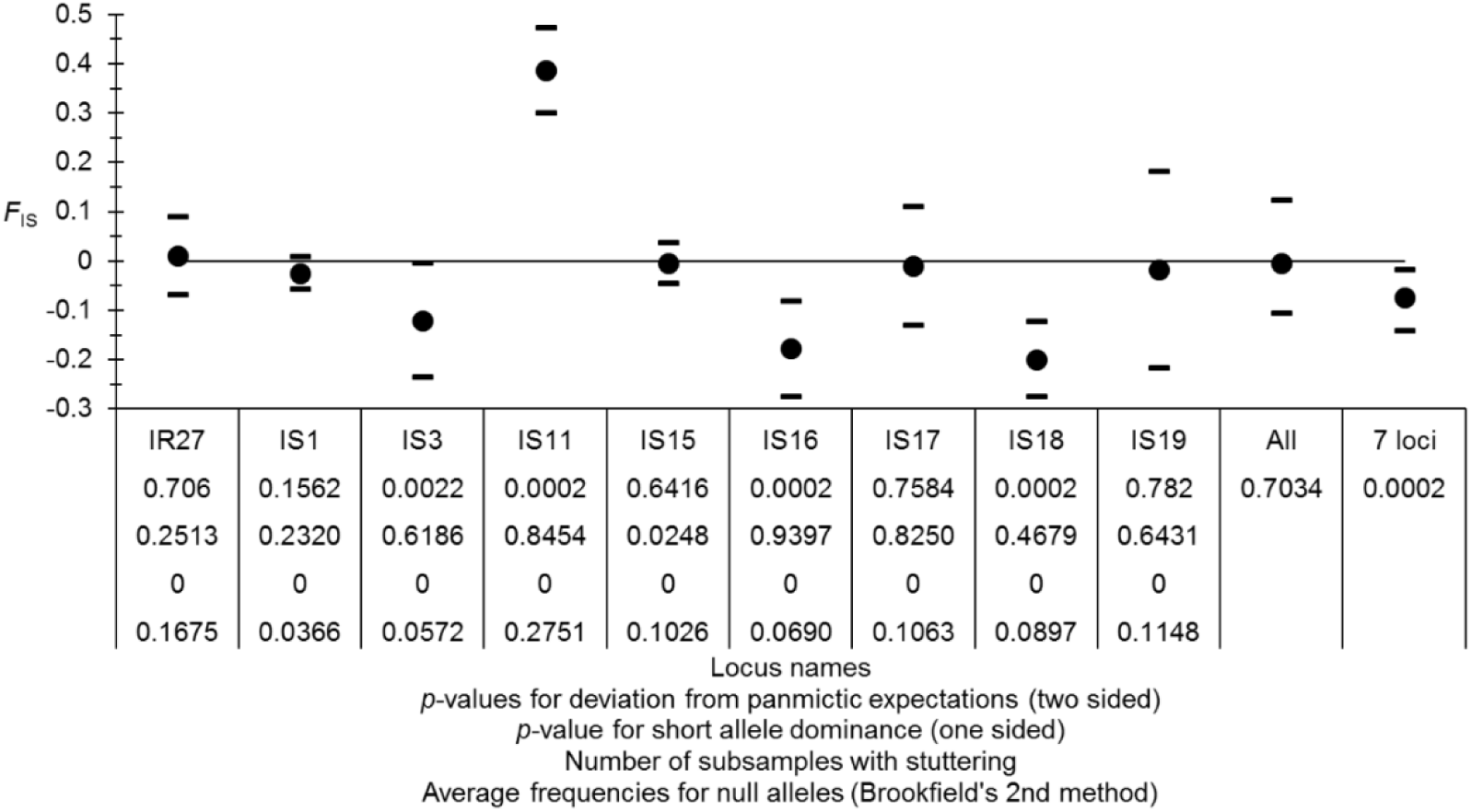
*F*_IS_ values for each locus, averaged across those (All), or over 7 loci without amplification problems (IS11 and IS15) for *Ixodes scapularis* cured data from the eastern U.S.A. with 95% jackknife confidence intervals over subsamples (for each locus) and bootstraps over loci (All). Results of tests for panmixia, short allele dominance, number of subsamples with stuttering for each locus, and null allele frequencies are also indicated.

**Figure 6.**
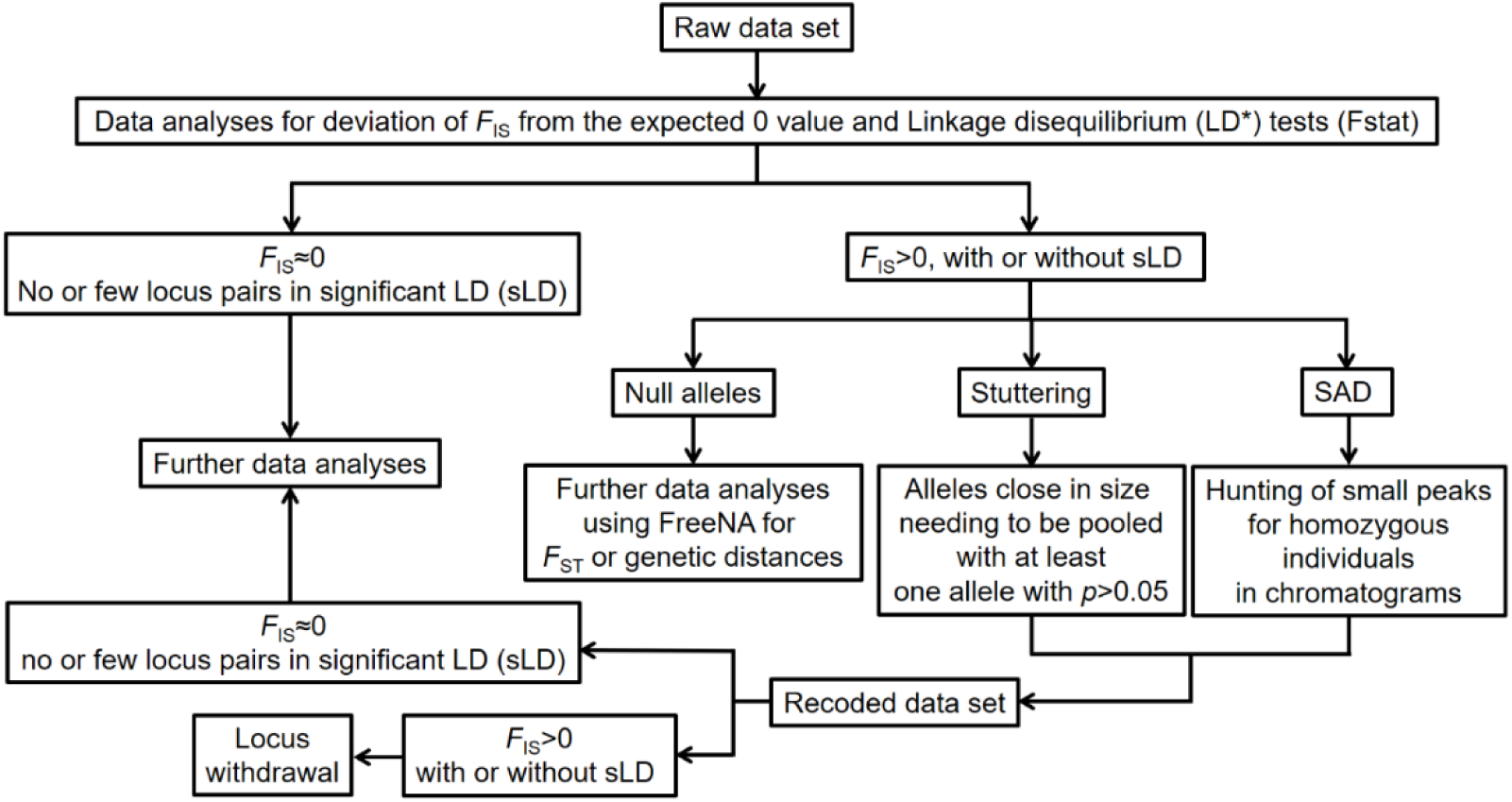
Flowchart of the recommended steps in population genetics data analyses.

Two loci displayed highly significant SAD (Figure 2): loci IS1 (*ρ*=-0.57, *p*-value=0.0015) and IS15 (*ρ*=-0.46, *p*-value=0.008), which stayed significant after BH correction (0.0135 and 0.036 respectively). Stuttering was diagnosed for four loci (IS1, IS15, IS16, IS17 and IS19). According to Brookfield’s second method, null allele frequencies could range between 0.06 and 0.28 on average (Figure 2), but these estimates do not correct for other errors and, as discussed above, null alleles only partly explain the observed *F*_IS_ and its variation across loci.

These heterozygote deficits and high proportion of significant LD may come from amplification problems detected as SAD, stuttering and null alleles. Amplification problems are expected to occur in individuals presenting an apostate DNA. This can happen if these individuals belong to lineages that are significantly divergent from the majority; when their DNA is partly degraded, at low concentration; or a combination of these different causes. This may amplify a preexisting correlation between allele occurrences at different loci. One way to test for this is to study the correlation between the number of missing genotypes (blanks) of an individual and its number of heterozygous sites. This correlation is expected to be negative if amplification problems occur more often in some individuals than in others (Kaboré et al., 2011). A Spearman’s rank correlation test with Rcmdr outputted *ρ*=-0.26 (*p*-value<0.0001).

### Cured data set

To prevent the omission of five (out of nine) loci, due to stuttering, SAD and/or possible selection, we went back to the data. We first scanned the chromatograms for previously ignored micro-peaks in homozygous individuals at loci displaying SAD (IS1 and IS15) (Figure 4). For IS1 and IS15, SAD might well have explained why we also found stuttering at these locus (see below). We then tried to pool alleles close in size as described above for loci IS16, IS17 and IS19. For IS16, alleles 88, 90 and 92 were recoded as 94; and allele 96 as 98. For locus IS17, alleles 170 and 172 were recoded as 174; alleles 178, 180, 182 and 184 were recoded as 186; alleles 188, 190 and 192 were recoded as 194; and alleles 196 and 198 as 200. Finally, for locus IS19, allele 91 was recoded as 93; alleles 97, 99, 101 and 103 as105; and alleles 107 and 109 as 111. The obtained amended dataset was called “Cured dataset” (Supplementary file S1). Pooling alleles only increases homoplasy. Everything being equal, the effect of homoplasy on *F*_IS_ or *F*_ST_ is equivalent to a mutation rate increase by a factor *K*/(*K*-1) (Rousset, 1996), where *K* is the number of possible alleles. Thus, for microsatellite loci with many possible alleles, the effect is deemed negligible, especially for *F*_IS_. Here, the resulting number of alleles per locus was 17 on average, with 5 and 27 as the outermost-lying values. The resulting homoplasy effect on *F*-statistics can thus be safely ignored.

With the cured dataset, the proportion of locus pairs in significant LD dropped to 19%, but more importantly, the smallest *p*-value after BY correction was 0.0797. The correlation between NLD and *H*_T_ became positive (*ρ*=0.57, *p*-value=0.1056). This change may point towards a Wahlund effect (Manangwa et al., 2019). Nevertheless, this would be incompatible with the *F*_IS_ observed (see below) and with the fact that none of these tests remained significant after BY adjustment. A heterozygote deficit was no longer observed (*F*_IS_=-0.004 in 95%CI=[-0.105, 0.124], bilateral *p*-value=0.7034) (Figure 5). StrdErrFIS=0.064 was three times higher than StrdErrFST=0.022. There was no correlation between *F*_IS_ and *F*_ST_ (*ρ*=-0.27, *p*-value=0.77). The correlation between number of missing genotypes and *F*_IS_ was positive, though marginally not significant (*ρ*=0.52, *p*-value=0.0809). MicroChecker diagnosed null alleles in 11 subsamples for locus IS11 and in one subsample for loci IR39 and IS19, which correspondingly displayed relatively high and variable *F*_IS_ and substantial proportions of missing genotypes (see above and Figure 5). IS15 still displayed SAD (*ρ*=-0.3814, *p*-value=0.0248), though with much less intensity. In addition, after BH correction, the test was not significant any longer (*p*-value=0.2232). When IS11 and IS15 were removed from the dataset, global *F*_IS_ became significantly negative (*F*_IS_=-0.074 in 95%CI=[-0.142, - 0.017], *p*-value=0.0002) as expected for small subpopulations in dioecious species (Balloux, 2004). As for subdivision, *F*_ST_ remained almost unaffected, even with the ENA method (*F*_ST_=0.103 in 95%CI=[0.067..0.14], *p*-value<0.0001).

### Conclusion

Combinations of amplification errors manifesting as null alleles, SAD or stuttering lead not only to heterozygote deficits, but an overall increase in LD. In order to correct for SAD, it may be useful to hunt for small and hard-to-detect peaks in chromatograms for homozygous individuals and pool alleles close in size to correct for stuttering. Uncorrected, these problems have the potential to lead to the unnecessary withdrawal of the “flawed” loci. These proposed amendments, together with null allele management with the ENA algorithm (Chapuis and Estoup, 2007), resulted in analyses revealing heterozygote excesses, as expected in small dioecious subpopulations. These changes also reduced the proportion of significant LD tests which, more importantly, were no longer significant after BY correction. Such corrections can then lead to more accurate estimates of the degree of population subdivision, as shown elsewhere for null alleles (Chapuis and Estoup, 2007; Séré et al., 2017b). The different steps to be followed during population genetics data analyses of such datasets are summarized as a flowchart in Figure 6.

In our case, the relatively important global LD across loci is probably due to small effective sizes of the *I. scapularis* subpopulations (Waples, 2006; Waples and Do, 2010). Additionally, null alleles are still influencing the cured data, and predominantly affect individuals that display some kind of deviating DNA (as explained in the introduction) and may also contribute to inflate the perceived LD. In fact, the correlation between the number of missing genotypes and the number of heterozygous sites was still significantly negative in the cured data set (*ρ*=-0.32, *p*-value<0.0001).

It is worthy to note that a reanalysis of the raw data set using MicroDrop 1.01 (Wang et al., 2012) resulted in smaller *F*_IS_ and *F*_ST_ values and only three significant LD tests, none of which stayed significant after BY correction (though the smaller *p*-value=0.0556 was marginal). However, locus IS15 still displayed a significant SAD, (*p*-value=0.0085, and corrected *p*-value=0.0765) and stuttering still was detected once, twice and three times for loci IS1, IS16 and IS17, respectively, and appeared once for locus IR27. MicroDrop takes into consideration that heterozygote deficits and missing data are entirely due to allelic dropouts. It is difficult to thoroughly understand these results since here we have convincing evidence that the amplification problems were due to SAD, stuttering and null alleles. This issue would require a full simulation study with different scenarios and a comparison of methods. That MicroDrop may efficiently cure not only allelic dropouts but any kind of amplification problem without any bias offers the potential for an extremely valuable tool for the scientific community, however, this remains to be ascertained. Nevertheless, MicroDrop did not entirely cure the data from SAD and stuttering, which does not advocate for the efficacy of the algorithm used in this software. As far as our *I. scapularis* data set is concerned, the cures proposed provided satisfactory results and additional useful tools to those already proposed in recently published papers on the detection and identification of causes of heterozygote deficits (Waples, 2015; De Meeûs, 2018b; Waples, 2018; Manangwa et al., 2019).

Subdivision into small and isolated subpopulations was confirmed by the relatively small effective population size estimated from *F*_IS_ (without IS11 and IS15) with Balloux’s method (Balloux, 2004) (*N*_*e*_≈7 in 95%CI=[4, 29] individuals), and the relatively important subdivision measurements between contemporaneous subsamples (to avoid temporal effects) in cohorts where these were possible (i.e. cohorts 5, 7, 8 and 9) (average *F*_ST_’≈0.19 in 95%CI=[0.1, 0.29]). This provided an estimate for an immigration rate of *Nm*≈1 individual per generation. These results contrast with those obtained from the closely related European *I. ricinus*, with no or weak LD in adults, displaying Wahlund effects at small scales, and with much weaker subdivisions (Delaye et al., 1997; De Meeûs et al., 2002; Kempf et al., 2010). The exact localization of subsamples was known. Consequently, there was no real need to undertake a Bayesian clustering approach as frequently met in publications. We nevertheless undertook DAPC analyses (Jombart et al., 2010) on the cured data set, which provided strong partitions (mean assignment from 0.96 to 1), but different from one run to the other and a poor correlation with geographic coordinates (Figure 7). There are indeed many instances where the relevance of such approaches may be debatable (Latch et al., 2006; Kaeuffer et al., 2007; Frantz et al., 2009; Meirmans, 2015; Manangwa et al., 2019). By contrast, with regard to the aim of this study, clustering techniques are useful to detect a Wahlund effect. Structure (and other software) can be very helpful to estimate the race or species assignment of different individuals of a population, but this was not the aim of the study. The fact that we obtain, with the cured data set, substantially negative *F*_IS_ and substantially high *F*_ST_ estimates obviously argues in favor of strong population subdivision. The estimates of immigration and effective subpopulation sizes (here *N*_*e*_*m*=1 and *N*_*e*_=7) illustrate this point and support the idea that this tick population is strongly subdivided.

**Figure 7:**
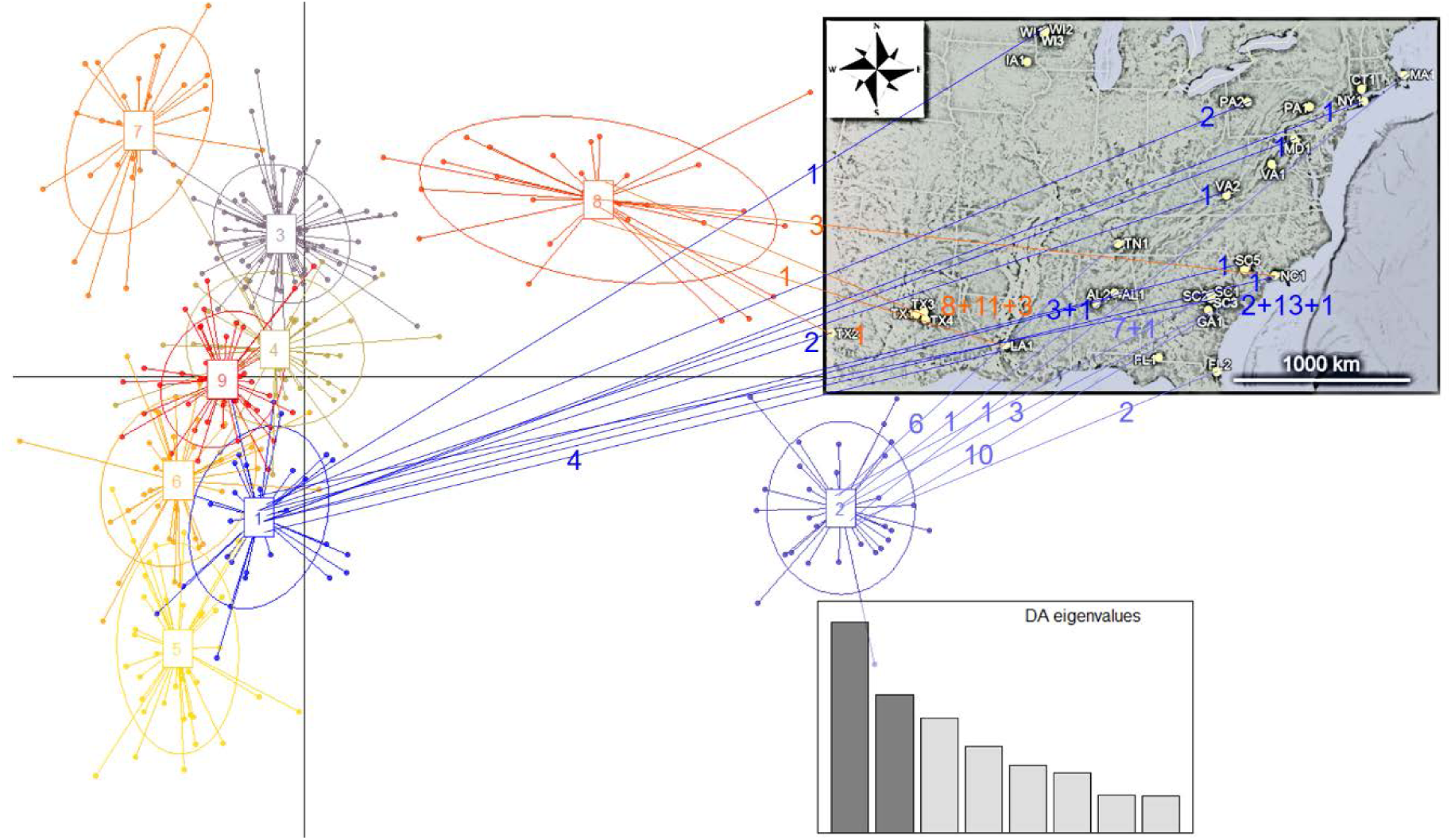
DAPC results with the maximum number of clusters set to 10, which resulted in 9 clusters. Individuals’ original sites and their numbers are indicated with the same color of the cluster to which they belong for clusters 1 (blue), 2 (violet-blue) and 8 (orange). Even if some geographic concordance can be noticed (e.g. Cluster 8 is mainly in Texas), many individuals that belong to the same cluster originated from distant sites. Changing the maximum number of clusters (e.g. 100) provided other optimal number of clusters (e.g. 7) but with similar characteristics as the one observed here.

Finally, we still do not know if mitochondrial clades have any real biological meaning for *I. scapularis* population structure or systematics. This will be treated in detail in a further study that will also include immature stages.

## Data accessibility

The raw and cured datasets are available as “supplementary file S1” at: http://www.t-de-meeus.fr/Data/DeMeeus-et-al-SAD&StutteringI-scapularisUSA-PCI-EvolBiol-TableS1.xlsx

## Supplementary material

Script and codes for the DAPC analysis are in Appendix 1

## Acknowledgements

This work was funded by NSF Grant # EF0914390 to L.B. and EEID EF-0914476 to J.T. We thank the members of the Lyme Gradient Consortium and many individuals who provided ticks. We are also grateful to Heather Walker, Jenny Dickson, Keely Duff, Laquita Burton, Alysha Benn and Nina Griffin, and the several other undergraduate students who provided field and laboratory assistance. We would like to thank one anonymous referee, Martin Husemann and Eric Petit for their comments and suggestions that helped with improving the present manuscript. This preprint has been peer-reviewed and recommended by Peer Community In Evolutionary Biology (https://doi.org/10.24072/pci.evolbiol.100081)

## Conflict of interest disclosure

The authors of this preprint declare that they have no financial conflict of interest with the content of this article. TdM is recommender for PCI Evol Biol and PCI Ecology

## Appendix

### Appendix 1: Script for the DAPC analyses with 100 or 10 as maximum possible number of clusters

~~~
>IscapCuredAll<-read.table(“IscapCuredDataDAPC.txt”, header=TRUE,
sep=“\t”, na.strings=“NA”, dec=“·”, strip.white=TRUE)
> IscapAdeCuredAll<-df2genind(IscapCuredAll, sep = NULL, ncode =
3, ind.names = NULL, loc.names = NULL, pop = NULL, NA.char = “NA”,
ploidy = 2, type = “codom”, strata = NULL, hierarchy = NULL)
> x<-IscapAdeCuredAll
> grp<-find.clusters(x,max.n.clust=10)
Choose the number PCs to retain (>= 1): 200
Choose the number of clusters (>=2: 9
> dapc2 <- dapc(x, grp$grp, n.pca= NULL, n.da= NULL, var.contrib =
TRUE, scale = FALSE)
Choose the number PCs to retain (>=1): 200
Choose the number discriminant functions to retain (>=1): 8
> scatter(dapc2)
> summary(dapc2)
$n.dim
[1] 8
$n.pop
[1] 9
$assign.prop
[1] 0.9922481
$assign.per.pop
      1       2       3       4        5           6
7       8       9
1.0000000 1.0000000 1.0000000 0.9772727 1.0000000 0.9767442
1.0000000 1.0000000 0.9811321
$prior.grp.size
1   2  3  4  5  6  7  8  9
34 31 85 44 42 43 28 27 53
$post.grp.size
 1  2  3  4  5  6  7  8  9
34 31 86 44 42 42 29 27 52
>tabIscapCuredAllMaxClust109Clust<-
data.frame(Cluster=c(grp$grp),Proportion_assign_cluster
=dapc1$posterior, geno=IscapCuredAll)
>
write.table(tabIscapCuredAllMaxClust109Clust,”IscapCuredAllMaxClus
t10-9Clust.txt”,col=NA, sep=“\t”, dec=“.”)
~~~

## Notes

http://www.t-de-meeus.fr/Data/DeMeeus-et-al-SAD&StutteringI-scapularisUSA-PCI-EvolBiol-TableS1.xlsx

